# Atomic Structure and Phospholipid Binding Properties of the *Francisella* Type VI Secretion System Effector Protein PdpC

**DOI:** 10.1101/2025.05.08.652963

**Authors:** Xiaoyu Liu, Daniel L. Clemens, Bai-Yu Lee, Xian Xia, Hongcheng Fan, Kaelyn Y. Feng, Marcus A. Horwitz, Z. Hong Zhou

**Affiliations:** Department of Microbiology, Immunology and Molecular Genetics, University of California, Los Angeles (UCLA), Los Angeles, CA 90095, USA; The California NanoSystems Institute (CNSI), UCLA, Los Angeles, CA 90095, USA; Department of Medicine, UCLA, Los Angeles, CA 90095, USA

**Author notes:** Correspondence (cryoEM and structures) and (infectious diseases and biochemistry). Contributed equally (Xiaoyu Liu, Daniel L. Clemens, Bai-Yu Lee).

**Keywords:** *Francisella*, Type VI Secretion System, Effector Protein, cryoEM, Phospholipid binding, PI3P

## Abstract

*Francisella tularensis*, a gram-negative facultative intracellular pathogen, causes tularemia, a potentially fatal infection. The *Francisella* pathogenicity island (FPI) encodes a type VI secretion system (T6SS), critical for the bacterium’s ability to replicate within host cells and cause disease. PdpC, an FPI-encoded T6SS effector protein, is essential for phagosomal escape, intracellular replication, and virulence in animals, yet its structure and function remain poorly understood. Here, we expressed PdpC recombinantly and determined the cryoEM structure of PdpC at 3.4 Å resolution, revealing a monomeric 156 kDa protein with a seahorse-shaped architecture (80 Å × 75 Å × 120 Å). PdpC comprises five domains: an N-terminal domain (NTD) forming a prominent lateral head lobe, a central body domain, a C-terminal tail domain, a small wedge domain at the groove between the body and tail, and an unmodeled “mouth” domain that completes the seahorse-like shape. Functional studies demonstrate that PdpC binds specific host cell phospholipids, including phosphatidylinositol-3-phosphate. These findings provide a foundation for elucidating PdpC’s mechanism of action and developing strategies to prevent or treat tularemia.

## INTRODUCTION

*Francisella tularensis* (Ft) subsp. *tularensis* is a gram-negative facultative intracellular bacterial pathogen that causes a serious and potentially fatal infection, tularemia, in humans^1, 2^. Because of its high infectivity and capacity to cause severe morbidity and mortality, Ft subsp. *tularensis* is considered an agent of bioterrorism and is classified as a Tier 1 select agent. Macrophages are the primary host cells for *F. tularensis* in animals^3^. After uptake by macrophages via the novel mechanism of looping phagocytosis^4^, *F. tularensis* initially resides in a phagosome that resists fusion with lysosomes and fails to acquire acid hydrolases, such as cathepsin D^5^. Within hours, the bacterium escapes from its phagosome and multiplies extensively in the macrophage cytosol^6^.

*F. novicida* (Fn) (also known as *F. tularensis* subsp. *novicida*) is often used as a model for Ft subsp. *tularensis* because it is genetically closely related and exhibits the same intracellular lifestyle but is less virulent to humans. In addition, whereas Ft has two copies of its FPI, Fn has only a single copy, thereby facilitating genetic manipulations. The *Francisella* pathogenicity island (FPI) is essential to the capacity of both Ft and Fn to escape from their phagosome, replicate inside macrophages, and cause fatal disease in animals^7, 8^. The FPI encodes a type VI secretion system (T6SS) that secretes effector molecules that act on the target cell. Several proteins encoded within the FPI have been shown to be secreted in a T6SS- dependent fashion, including IglC, VgrG, PdpA, IglG, PdpC, and PdpD^9–11^. Four additional proteins encoded outside of the FPI (OpiA and OpiB1-OpiB3) also have been shown to be secreted in a T6SS-dependent fashion^11^. AnmK, encoded within the FPI, is thought to be a T6SS secreted anhydro-N-acetylmuramic acid kinase, but its T6SS-dependent secretion has not been verified and its role in pathogenesis is unclear. It is unclear whether VgrG, PdpA, IglC, and IglG, which are essential for T6SS-mediated protein secretion, solely serve as structural components secreted alongside effectors or also exhibit effector functions in infected host cells. On the other hand, disruption of genes encoding PdpC, PdpD, OpiA, or OpiB1 – OpiB3 does not impact secretion of other T6SS proteins^11, 12^, supporting their role as secreted effector proteins. Deletion of the *pdpC* gene in *F. novicida* has been shown to cause a severe defect in phagosome escape and a partial defect in virulence in mice^13^. Deletion of both *pdpC* and *pdpD* causes an even more profound defect in *F. novicida* phagosome escape in macrophages and combined deletion of *pdpC*, *pdpD*, and *anmK* renders *F. novicida* avirulent in a mouse model^13^. In the *Galleria mellonella* (wax moth) larva model, the presence of an individual gene for *pdpC*, *pdpD*, or *opiA* alone, together with functional T6SS secretory capacity, is sufficient for full virulence^12^. In the SCHU strain of *F. tularensis* subsp. *tularensis*, PdpC has been shown to be essential to the capacity of the bacteria to grow in macrophages and cause disease in mice^14^ and an intermediate phenotype with decreased phagosome escape in macrophages and attenuated virulence in mice has been shown for the Schu S4 strain with deletion of both *pdpC* genes^15^. While OpiA has been demonstrated to be a PI3-kinase that delays phagosome maturation^16^, the biological functions of other secreted effector proteins remain largely unknown.

The current study aims to elucidate the structure and function of PdpC, which is one of the key effector proteins encoded by the FPI and secreted via the T6SS. Although PdpC has been implicated in phagosomal escape and virulence, its molecular architecture and mechanistic role remain poorly understood. Using cryoEM, we determined the atomic structure of recombinantly expressed *Francisella* PdpC at 3.4 Å resolution, revealing a seahorse-like monomeric architecture composed of five domains. We further investigated its biochemical properties and found that PdpC binds selectively to host phospholipids, including phosphatidylinositol-3-phosphate. These findings establish a structural and functional framework for understanding the role of PdpC in *Francisella* pathogenesis.

## RESULTS

### Expression and isolation of PdpC from *F. novicida*

PdpC is a key effector protein secreted via T6SS (Fig. 1A). To facilitate purification of PdpC, we prepared *F. novicida* expressing N-terminal His6-FLAG epitope tagged PdpC. However, we found that samples purified from this strain using nickel-agarose affinity chromatography and gel filtration were contaminated with FPM13 (FTN_1118) due to its metal binding properties, similar native molecular weight and overall abundance (manuscript submitted, see BIORXIV/2025/652791). Therefore, we prepared *F. novicida* expressing N-terminal His6-FLAG epitope tagged PdpC on a ΔFPM13 background. This strain was inoculated into 3 liters of Trypticase Soy Broth with 0.2% cysteine (TSBC), 5 mM betaine, 0.1 mg/mL FeSO4, 0.2 mg/ml hygromycin, containing 5% KCl to induce formation of the T6SS^9^. The bacteria were lysed by sonication and PdpC was purified by Ni-agarose and gel filtration chromatography (Fig. 1B). On gel filtration chromatography, the fractions with the most intense staining by Sypro Ruby for a 150 kDa band (fractions 24 – 26, Fig. 1B) correspond to the elution position of the gel filtration calibration standard, alcohol dehydrogenase. The protein in fractions 24 – 26 were concentrated with a 100 kDa MWCO centrifugal concentrator (Millipore) and the buffer changed to 20 mM HEPES, 0.3 M NaCl, 3% glycerol, 0.1% CHAPS using a 40 kDa MWCO Zeba Spin Desalting column (Thermo Fisher).

**Fig 1.**
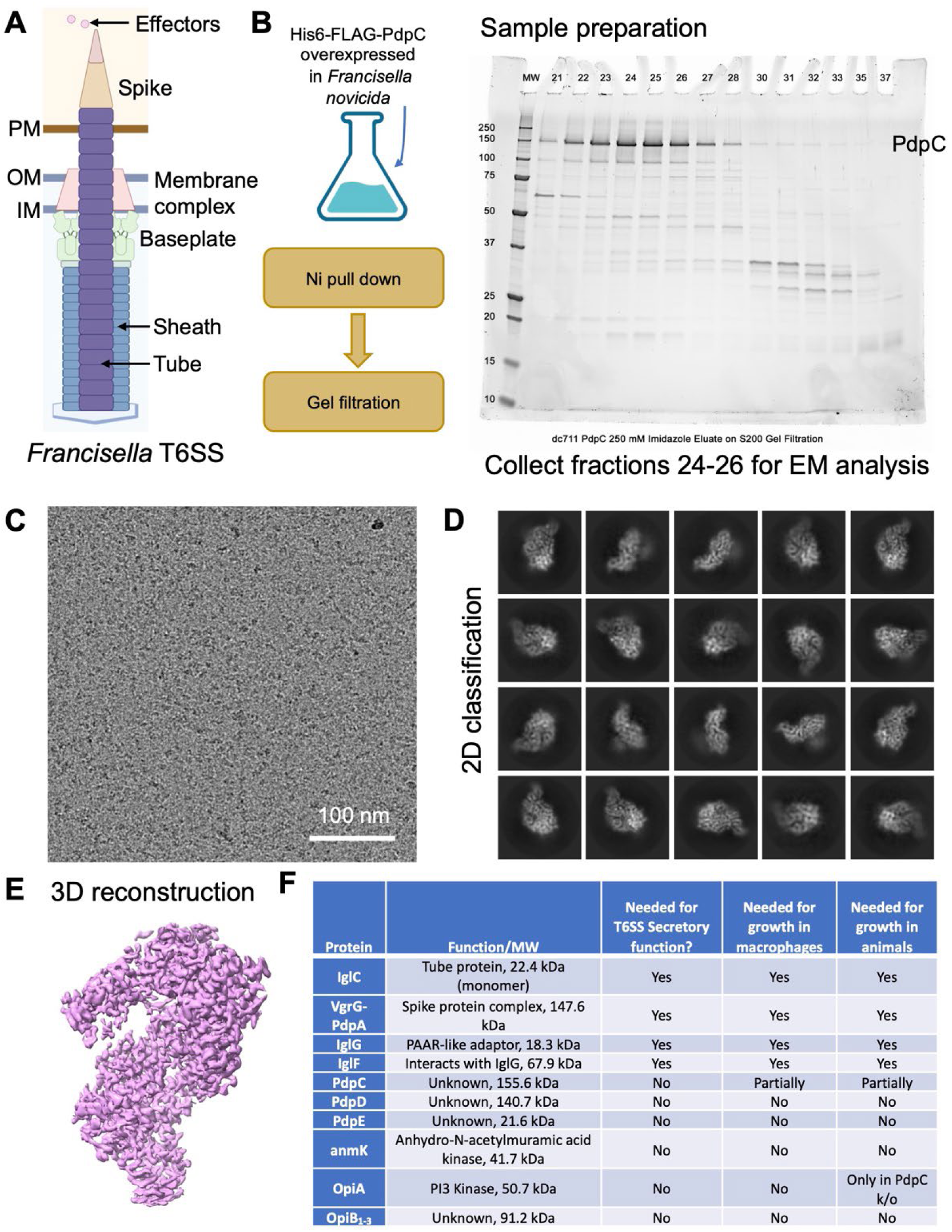
Structure determination of PdpC. **(A)** Schematic of *Francisella* T6SS. IM, OM, and PM refer to the inner membrane, outer membrane, and phagosomal membrane, respectively. **(B)** Protein sample preparation flowchart and the SDS-PAGE gel of PdpC. **(C)** CryoEM micrograph of PdpC. **(D)** Representative 2D class averages of PdpC particles. **(E)** CryoEM reconstruction of PdpC. **(F)** *Francisella* proteins secreted via the T6SS apparatus.

### Determination of the atomic structure of PdpC

As mentioned above, in the initial attempts to determine the structure of PdpC, only donut-shaped particles were visible in the cryoEM images. Subsequent data processing, 3D reconstruction, and protein identification revealed that these particles corresponding to the FPM13 complex. When PdpC was overexpressed in a ΔFPM13 *F. novicida* strain, the donut-shaped particles were no longer observed. However, this led to a new challenge: no discernible particles could be seen on Quantifoil holy grids under the microscope. Extensive optimization was performed to overcome this, including adjusting the protein buffer composition, testing various grid types, and adding detergents to alter the properties of the air-water interface. Ultimately, graphene oxide (GO)-coated grids enabled the reconstruction reported in this study. CryoEM imaging and 2D classification revealed comma-shaped particles (Fig. 1C-D). Due to the low signal-to-noise ratio and severe preferred orientation of the particles, we employed spIsoNet^17^ during data processing to improve 3D reconstruction quality. This approach yielded a 3.4 Å resolution map (Fig. 1E), which supported atomic model building.

PdpC is a 156 kD protein consisting of 1325 amino acids and exits as a monomer in the reconstructed structure. No homologous structures of PdpC have been reported to date.

Moreover, the AlphaFold prediction of PdpC is of very low confidence and differs markedly from our cryoEM determined structure (Suppl. Fig. S2), posing additional challenges to model building. We initially used Model-Angelo^18^ to generate a starting model. Due to discontinuities in the cryoEM density in certain regions, the model was manually corrected and adjusted in Coot, followed by further refinement in Phenix^19^. This optimized workflow provides a useful reference for the structural determination of other T6SS effector proteins that are critical for *Francisella* replication and virulence (Fig. 1F).

### Domain organization and inter-domain interactions of PdpC

The cryoEM structure reveals that PdpC exists as a monomer in solution and adapts an asymmetric elongated shape, like a seahorse (Fig. 1E). It measures approximately 120 Å in height and 80 Å in width (Fig. 2A-B). The PdpC structure is primarily composed of α-helices, with the exception of the N-terminal domain (NTD), which adopts a compact immunoglobulin (Ig)-like β-sandwich fold (Fig. 2C-D). It can be divided into five domains arranged sequentially from the N- to C-terminus: the NTD (residues 25–169), which forms a prominent lateral head lobe; a central body domain (residues 209–787); an unmodeled region (residues 787-1027), referred to here as the “mouth” domain due to its location; a wedge domain (residues1027–1145), which bridges the body and tail domains; and a C-terminal tail domain (residues 1145– 1325), forming the tip of the molecule and completing the seahorse-like shape. While most of the structure was well resolved, a central ∼200-residue region corresponding to the mouth domain lacked interpretable density, likely due to conformational flexibility, and could not be modeled. This flexible segment is surrounded by the curved body, wedge, and tail domains, with the NTD extending laterally from the main structure. Importantly, structural homology searches revealed no similarity to any known protein in the PDB. These findings indicate that PdpC represents a novel protein fold of potential mechanistic and evolutionary significance.

**Fig 2.**
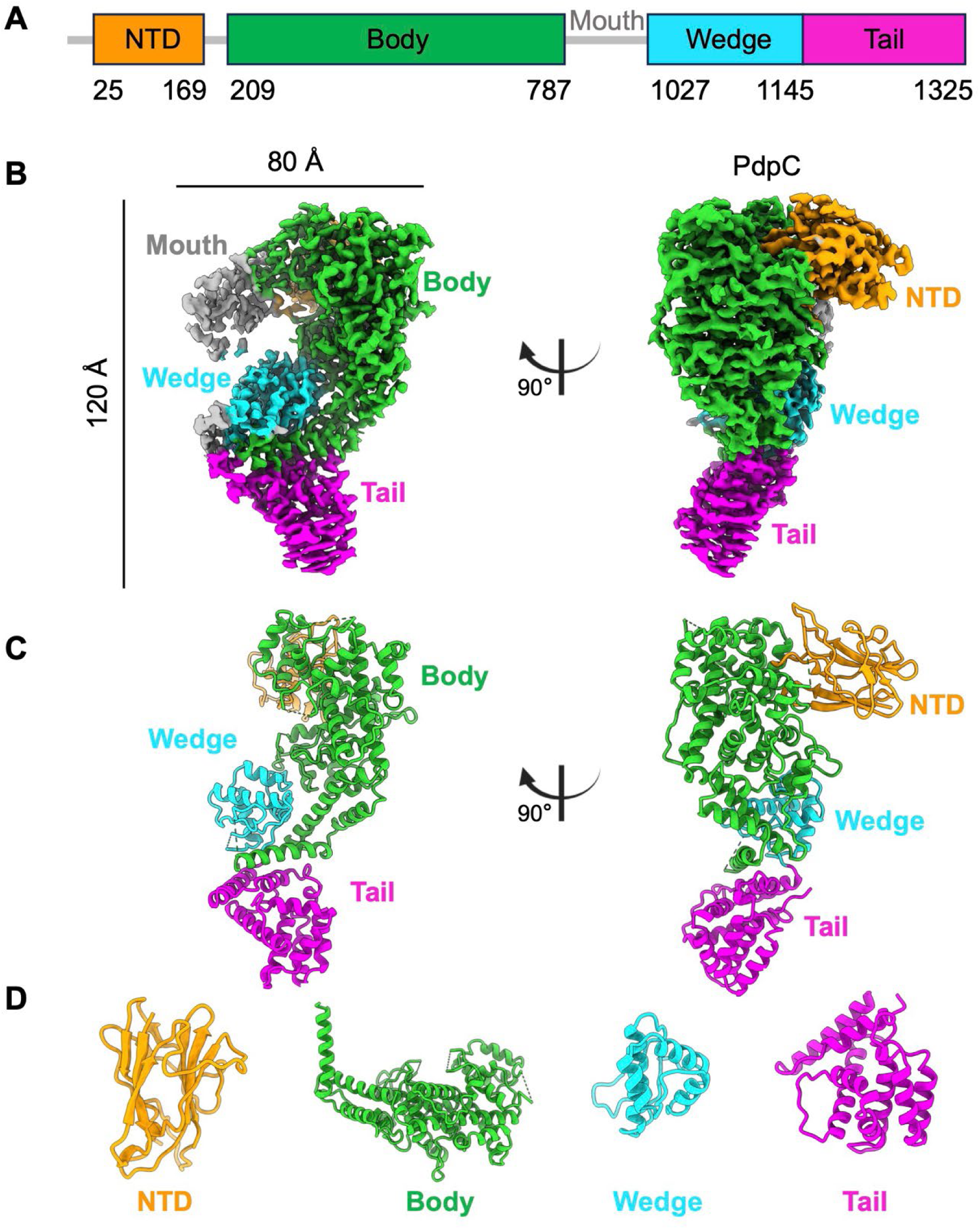
CryoEM structure of PdpC. **(A)** Domain organization of *Francisella* PdpC. Domains are labeled with different colors; unmodeled regions are shown in gray. **(B)** Two different views of the cryoEM density map of PdpC, with domains colored according to scheme in A. **(C)** Two different views of the PdpC structure, colored as in B. **(D)** Ribbon representation of individual domains.

Detailed analysis of the individual domains further reveal the structural uniqueness of PdpC (Fig. 2D). The NTD forms an Ig-like β-sandwich, composed of two β-sheets, and projects laterally from the main scaffold. The body domain, the largest of the five, forms the structural scaffold of the protein, consisting entirely of α-helices arranged in a helical solenoid-like fold. It forms the “spine” of the seahorse-like shape, connecting all other domains and potentially contributing to the structural rigidity of the protein. The wedge domain, a compact α-helical module, nestles into a groove between the body and tail domains, serving a likely stabilizing architectural role. The C-terminal tail domain is composed of several tightly packed α-helices forming a small, globular structure that caps the extended molecule. Together, these features suggest that PdpC is a modular, uniquely folded effector with no known structural homologs, reflecting evolutionary adaptation to its specialized role in *Francisella*’s type VI secretion system.

### PdpC selectively binds phospholipids

To gain insight into the biological function of PdpC, we examined the staining pattern of epitope- tagged PdpC when incubated with formaldehyde-fixed permeabilized PMA-differentiated human macrophage-like THP-1 cells that had ingested blue fluorescent 1 µm latex beads overnight as a marker of mature phagosomes (Fig. 3). The overall staining pattern was suggestive of staining of cellular membranes (Fig. 3). Little to no staining was observed of the latex bead phagosomes, which was not unexpected, as *Francisella-*containing phagosomes are known to resist maturation.

**Fig. 3.**
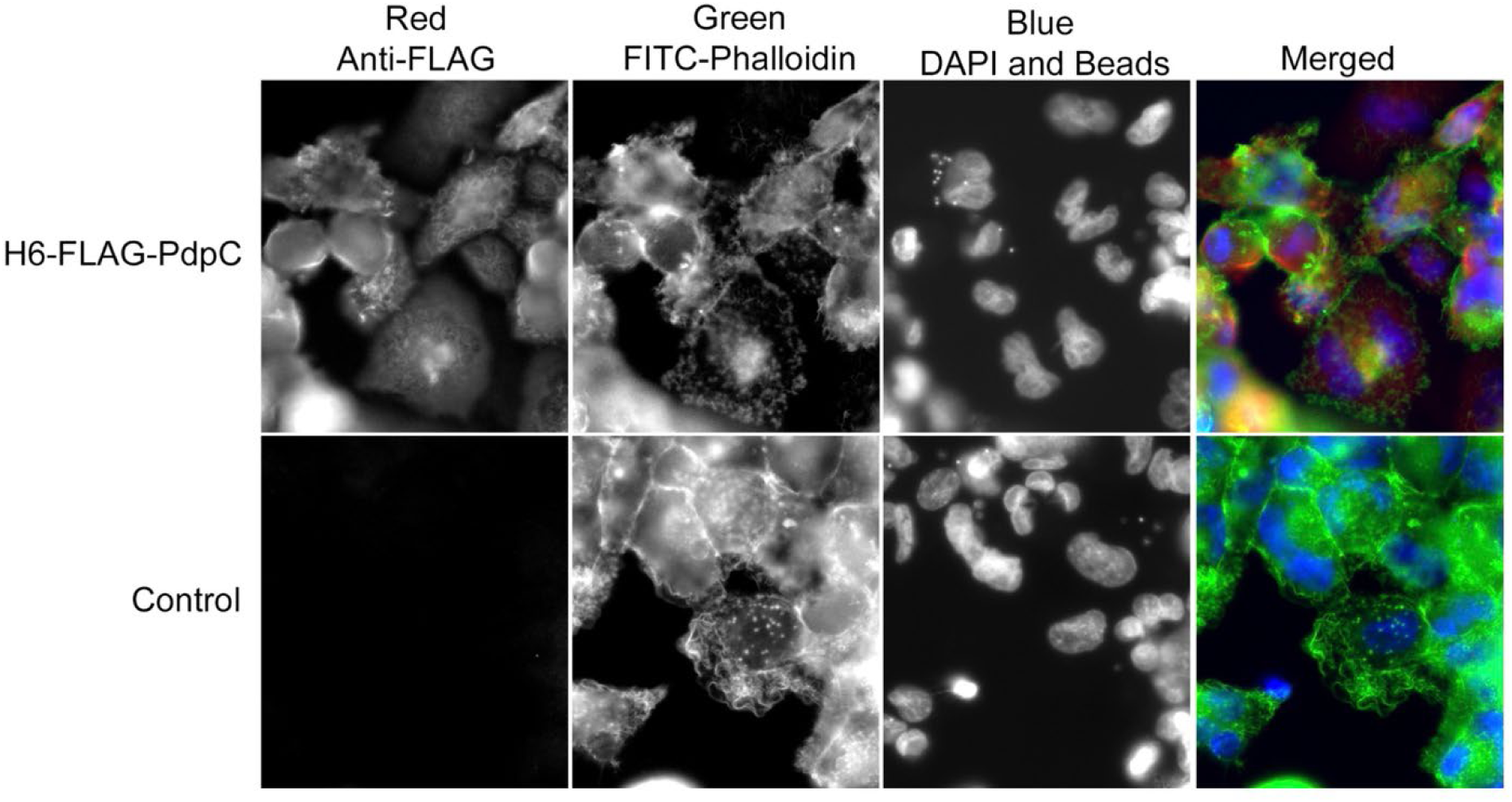
Staining of human macrophage-like THP-1 cells for PdpC. THP-1 cells were differentiated with PMA, allowed to uptake 1 micron blue fluorescent latex beads, fixed, permeabilized with 0.1% saponin, and incubated with 1 µg/mL H6-FLAG-PdpC in 0.1% TX-100, 1% bovine serum albumin in PBS with secondary staining using M2-anti-FLAG mouse antibody (1:300) and 1:50 Texas Red conjugated goat-anti-mouse IgG. Actin filaments were visualized by staining with FITC-phalloidin and nuclei were visualized by staining with DAPI.

Based on the observation of the immunofluorescent membrane staining pattern and the knowledge that many bacterial toxins and secreted effector proteins interact with host lipids, we next examined whether PdpC interacted with specific lipids by incubating purified FLAG-tagged PdpC with phospholipids on nitrocellulose strips (Echelon). Intense signals of FLAG-PdpC chemiluminescence were observed for PI(3)P, PI(3,5)P2, sulfatide and cardiolipin (Fig. 4). Much weaker signals were observed for PI(4)P, PI(3,4)P2, PI(4,5)P2, and PI(3,4,5)P3 (Fig. 4). Little to no signal was observed for phosphatidylinositol, cholesterol, phosphatidic acid, phosphatidylserine, phosphatidylethanolamine, phosphatidylcholine, triacylglycerol, diacylglycerol, sphingomyelin, and sphingosine-1-phosphate (Fig. 4).

**Fig. 4.**
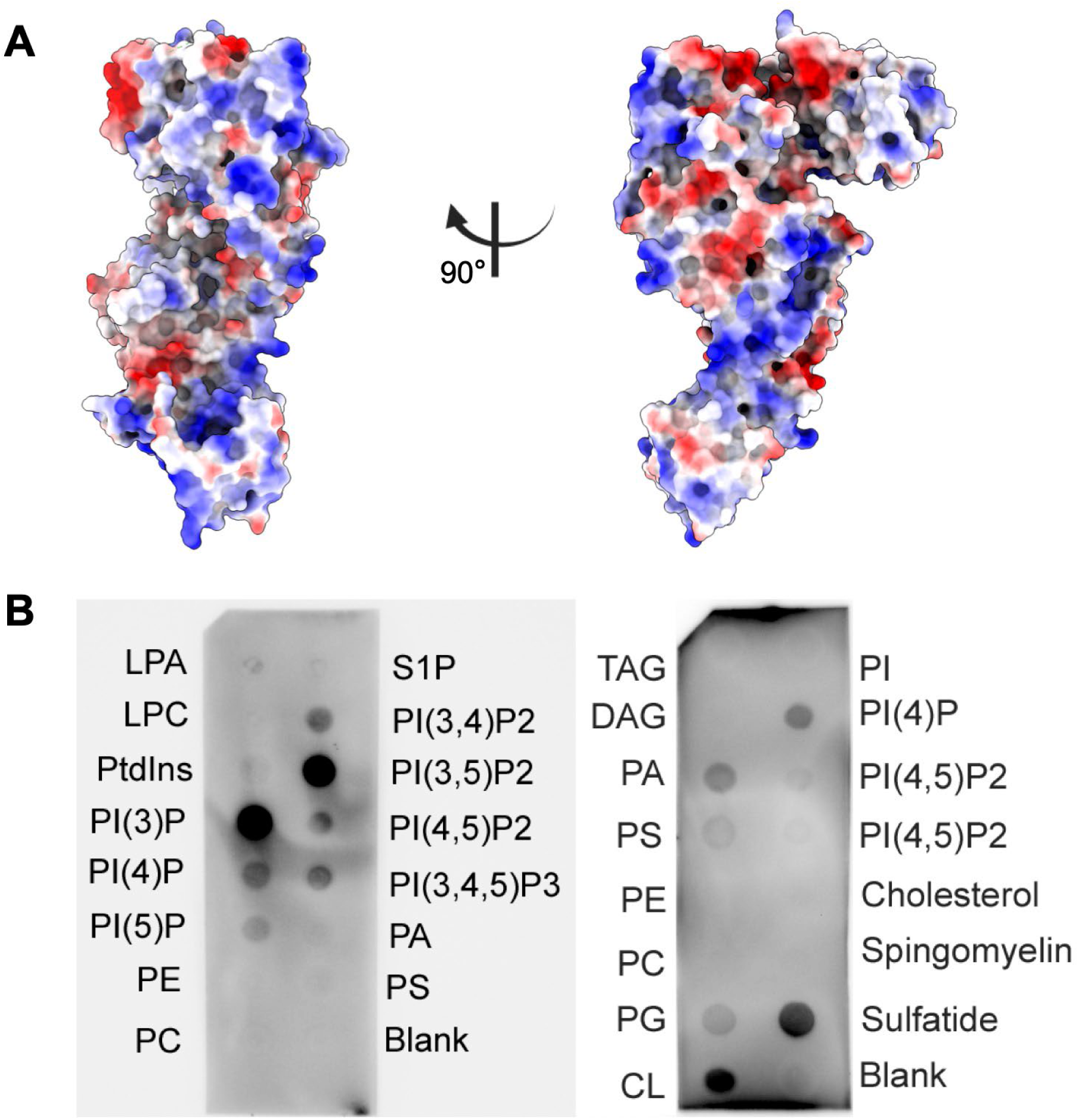
Phospholipid binding of PdpC. **(A)** Surface representation of the PdpC structure, with positive, neutral, and negative electrostatic potentials shown in blue, white and red, respectively. **(B)** Nitrocellulose strips with phospholipids (Echelon) were blocked by incubation with Tris buffered saline-BSA-Tween-20 and incubated with H6-FLAG-PdpC (5 μg/mL) in the blocking buffer, followed by monoclonal anti-FLAG-HRP and chemiluminescent substrate. Strong staining of PI(3)P and PI(3,5)P2 are shown in the panel on the left and strong staining of sulfatide and cardiolipin (CL) are shown on the right. LPA, lysophosphatidic acid. LPC, lysophosphatidylcholine, PtdIns, phosphatidylinositol. PE, phosphatidylethanolamine. PC, phosphatidylcholine, S1P, sphingosine-1-phosphate. PA, phosphatidic acid. PS, phosphatidylserine. TAG, triacylglycerol. DAG, diacylglycerol. PG, phosphatidylglycerol. PI, phosphatidylinositol.

Whereas OpiA has been shown to be a PI3-kinase, we have not detected phosphoinositide kinase or phosphatase activity in PdpC (Supplementary Fig. S3). It is possible that a kinase or phosphatase activity exists that requires different cofactors or assay conditions than those that we tested.

The selective binding of PdpC to PI(3)P, which is localized on the cytosolic face of early phagosomes, would enable PdpC to be anchored to the *Francisella* phagosome and thereby exert its actions locally rather than globally following secretion. The nature of its biological function remains to be determined but could include facilitating phagosome escape and/or retarding phagosome maturation. It is noteworthy that binding to phosphoinositides is a common strategy of bacterial toxins and effector proteins that have extremely diverse biological functions^20–24^.

## DISCUSSION

We have purified and determined the atomic structure of the *F. novicida* T6SS secreted effector protein PdpC. The protein has a novel architecture and structural fold as DALI search^25^ reveals no similar structures in the protein data bank. BLAST searches^26^ also did not reveal any similar proteins with sequence homology outside of the *Francisella* genus. Within the *Francisella* genus, PdpC orthologues are largely restricted to *F. tularensis* and *F. novicida* species. A phylogenetic analysis^27^ noted that while PdpC orthologues are present in almost all Clade A species of *Francisella* (pathogens of terrestrial animals: *Francisella tularensis* subspecies *tularensis*, *holarctica*, *mediasiatica*, and *novicida*) with very high degree of identity (94%-100%). PdpC is absent from almost all Clade B species (pathogens of aquatic animals: e.g., *F. philomiragia* and *F. salina*) as well as the diverse Clade C cluster associated with marine environments and air conditioning systems (e.g., *F. frigiditurris*, *F. endociliophora*, *F. halioticida*, and *F. uliginis*). The extent to which Clade B and Clade C species of *Francisella* share the lifestyle and virulence mechanisms of the Clade A species has not been reported.

While PdpC is important to phagosome escape and virulence, its mechanism of action has not been determined. OpiA has been shown to be a PI3-kinase that may ensure the presence of PI(3)P on the *F. novicida* phagosome. As PI(3)P is present on the cytosolic aspect of early endosomes and early phagosomes, the PI3-kinase activity of OpiA may help to ensure the continued presence of PI3P on the *Francisella* phagosome prior to phagosome disruption^16^. We hypothesize that the phospholipid binding activity of PdpC anchors it to the cytosolic side of the *Francisella* phagosome so that its biological effector activity is focused on the phagosome rather than exerted globally in the cell. It is noteworthy that numerous *Legionella pneumophila* T4SS secreted effectors bind PI(3)P or PI(4)P as an anchoring mechanism but have a variety of other effector activities^28^.

Our data show that PdpC also binds to PI(3,5)P2, sulfatide (3-sulfogalactosyl ceramide), and cardiolipin. Sulfatide has been shown to be present on early and recycling endosomes^29^.

PI(3,5)P2 is less abundant than other phosphoinositides, but is thought to be present on the cytosolic aspect of early and late endosomes and lysosomes and may play a role in regulation of phagosome maturation and autophagy^30^. Cardiolipin is primarily found on the inner leaflet of mitochondrial membranes, but in early phases of apoptosis relocates to the surface of mitochondria and other intracellular organelles^31^. Binding to lipids in these locations may help to focus PdpC effector action on phagosome maturation and autophagic functions within the infected host cell.

The primary sequence and atomic structure of many bacterial effectors, including *Francisella* PdpC, frequently fail to predict their pathophysiological functions, as noted in prior studies^32^. This challenge underscores the complexity of effector-mediated disease mechanisms, particularly in tularemia, where PdpC’s precise role remains elusive. While future discoveries of structurally analogous proteins may provide comparative insights, the current lack of such parallels necessitates rigorous investigations to elucidate PdpC’s biological function in *Francisella* pathogenesis. The present study marks a pivotal advance as the first systematic characterization of PdpC, determining its atomic structure and demonstrating its lipid-binding properties. These findings establish a critical molecular framework that will facilitate subsequent research efforts to uncover PdpC’s contributions to tularemia virulence. By revealing PdpC’s structural and biochemical properties, this work paves the way for functional studies, such as host-pathogen interaction assays or targeted mutagenesis, to map its role in immune modulation or cellular disruption. Furthermore, the structural data may guide the design of novel therapeutics, such as inhibitors targeting PdpC’s lipid interactions. Therefore, this study not only enhances our understanding of *Francisella* biology but also contributes to the broader field of bacterial effector research, offering a foundation for combating tularemia and related infectious diseases.

## METHODS

### Construction of His-FLAG-PdpC expression vector

The promoter sequence of bacterioferritin (Pbfr) was amplified by PCR using genomic DNA prepared from *F. tularensis* subsp. *holarctica* Live Vaccine Strain. DNA sequences for a histidine tag (HHHHHH) followed by a thrombin cleavage site (LVPRGS) and a FLAG tag (DYKDHDGDY KDHDIDYKDDDDK) followed by a TEV protease cleavage site (ENLYFQG) were added to the 3’-end of Pbfr sequence by overlap extension PCR to generate the Pbfr-His6-FLAG fragment.

The sequence corresponding to nucleotides 4 to 3978 of pdpC (FTN_1319, 1325 amino acids) was amplified using *F. novicida* U112 genomic DNA as template. The rrnBT1/T2 terminator sequence was amplified from pDONR207 SARS-CoV NSP1, a gift from Fritz Roth^33^ (Addgene Plasmid #141255). These three PCR fragments were treated with appropirate restriction enzymes and cloned into pMP633 vector^34^ in between the MluI and BamHI restriction sites and confirmed by nucleotide sequencing (Laragen). Oligonucleotide primers used in the PCR reactions (Suppl. Table S2) and the plasmid map (Suppl. Fig. S1) are provided as supplementary information.

### Expression and Purification of PdpC

*F. novicida* expressing N-terminal His6-FLAG epitope tagged PdpC was inoculated into 3 liters of Trypticase Soy Broth with 0.2% cysteine (TSBC), 5mM betaine, 0.1 mg/mL FeSO4, 0.2 mg/ml hygromycin, containing 5% KCl to induce formation of the T6SS^9^. The bacteria were grown at 37°C with shaking at 140 rpm to an optical density (540 nm) of 2, pelleted by centrifugation, resuspended in 100 mL of 20 mM HEPES, pH 7.5, 0.3 M NaCl, 1 mM EDTA, 3 mM MgCl2, 10 mM Imidazole, 0.8% CHAPS, and 10% glycerol (lysis buffer) containing 1 mM PMSF, 1 mM N-ethylmaleimide, and 1:100 dilution of Protease Inhibitor Cocktail (MedChem Express, HY- K0010). The bacteria were lysed by sonication with a probe tip sonicator (Ultrasonics W-375) while cooling on ice and the sonicate was clarified by ultracentrifugation at 44,400 g for 90 min.

The clarified supernatant fluid was added to 1 mL of Ni-agarose (Pierce High-Capacity Ni-IMAC Resin), rotated end-over-end overnight at 4°C, washed with lysis buffer, transferred to a column, and eluted with lysis buffer containing 20 mM, 50 mM, 100 mM, and 250 mM imidazole (10 mL each) with collection of 3 mL fractions. The eluted fractions were analyzed by SDS-PAGE and fractions containing PdpC were combined, concentrated with a 100,000 MWCO centrifugal filter concentrator (Amicon Ultra-15) and applied to a Sephacryl S200 gel filtration column that had been pre-equilibrated with 20 mM HEPES, pH 7.5, 1 mM EDTA, 0.3 M NaCl, 10% glycerol, 1% CHAPS. Fractions were collected and analyzed by SDS-PAGE with protein staining by Sypro Ruby. Fractions with strong band at 150 kDa were pooled, concentrated with a 100,000 MWCO centrifugal filter concentrator and changed to a buffer containing 20 mM HEPES, 0.3 M NaCl, 3% glycerol, 0.1% CHAPS using a 40 kDa MWCO Zeba Spin Desalting column (Thermo Fisher) for use in cryoEM analysis.

### CryoEM sample preparation and image collection

For cryoEM sample preparation, we applied 3 μL of purified protein sample to the graphene oxide (GO)-coated EM grids (Shuimu BioSciences). The grid was incubated for 10 s and blotted for 12 s with filter paper (Ted Pella) at force 10, and then flash-frozen in liquid ethane using an FEI Vitrobot Mark IV (Thermo Fisher Scientific) set to 8 °C temperature and 100% humidity.

CryoEM grids were loaded into a Titan Krios electron microscope (Thermo Fisher Scientific) operated at 300 kV, equipped with a Gatan K3 Summit direct electron detector and a GIF Quantum LS energy filter set to a slit width of 20 eV. Automated data collection was performed using SerialEM^35^. The total dose rate on the sample was set to ∼50 e^-^/Å^2^/movie at a pixel size of 1.1 Å. In total, 39,482 movies were collected.

### Single particle cryoEM reconstruction and atomic model building

Motion correction and defocus estimation were performed in Live CryoSPARC^36^, followed by subsequent data processing using cryoSPARC. Particles were picked with Topaz^37^, and after extensive 2D and 3D classification, 616,877 particles were selected. Heterogeneous refinement was performed to remove low-quality particles, resulting in 322,627 particles retained for further data processing. Homogeneous and non-uniform refinements in cryoSPARC^36^ yielded reconstructions with density distortions due to severe preferred orientation. To overcome this, we exported the 322,627 particles to RELION 5^38^, where a modified version of the Blush regularization algorithm, inspired by spIsoNet^17^, was applied to improve particle alignment accuracy. This optimized workflow ultimately produced a distortion-free cryoEM map of PdpC at ∼3.4 Å resolution.

We first used ModelAngelo^18^ to generate an initial model. The atomic model of PdpC was then manually built and refined in COOT^39^, followed by further refinement in Phenix^19^. Refinement statistics of the model are summarized in Table S1. Figures were generated using USCF ChimeraX^40^.

### Lipid binding assay

Nitrocellulose strips with phospholipids (Echelon) were blocked by incubation with Tris buffered saline with 1% BSA and 0.1% Tween-20 and incubated overnight at 4°C with His6-FLAG-PdpC (5 μg/mL) in the blocking buffer. Strips were washed three times in TBS containing 0.1% Tween- 20 and incubated with mouse monoclonal anti-FLAG-HRP (1:30,000). Chemiluminescent signals were developed by incubating with Clarity Western ECL substrates (Bio-Rad) and detected using a ChemiDoc Imaging System.

### Phosphoinositide kinase and phosphatase assay

Epitope tagged PdpC (0.2 mg/mL), purified as described above, was incubated with 50 µM BODIPY-fluorescent labeled PI, PI(3)P, or PI(3,5)P2 PIPES (Echelon Biosciences) in 0.1 M PIPES, pH 7.0, 2 mM MgCl2, 0.5 mM MnCl2, with or without 10 mM ATP, 10 mM MgCl2, for 90 minutes at 37°C in total volumes of 25 µL. Reactions were stopped by the addition of 5 volumes of acetone, and the samples were dried under vacuum. BODIPY-labeled lipids were extracted with CHCl3, spotted onto silica thin layer chromatography plates, developed in CHCl3:MeOH:Acetone:HOAc:Water (70:50:20:20:20), and imaged under blue light with a BioRad ChemiDoc Imaging System.

## Acknowledgements

This project was supported by National Institutes of Health grant AI151055 (MAH and ZHZ). We acknowledge the use of resources at the Electron Imaging Center for NanoSystems supported by US NIH (1S10OD018111) and the US National Science Foundation (DBI-1338135 and DMR- 1548924).

## AUTHOR CONTRIBUTIONS

Z.H.Z. and M.A.H. supervised the project. D.L.C. and B.-Y.L. prepared the bacterial strains and protein sample; X.L. made cryoEM grids, performed cryoEM imaging, and data processing with help from X.X.; H.F. performed the spIsoNet reconstruction; X.L. built the atomic model and illustrated the structures; D.L.C. and B.-Y.L. performed immunofluorescence and lipid binding experiments. K.F. helped with the figure illustration. X.L., D.L.C., B.-Y.L., M.A.H. and Z.H.Z. interpreted the data and wrote the manuscript; and all authors reviewed and approved the paper.

## Data availability

The cryoEM density map has been deposited in the Electron Microscopy Data Bank under accession code xxxx. The atomic coordinate has been deposited in the Protein Data Bank under accession code xxxx.

---

**Supplementary Fig. S1:**
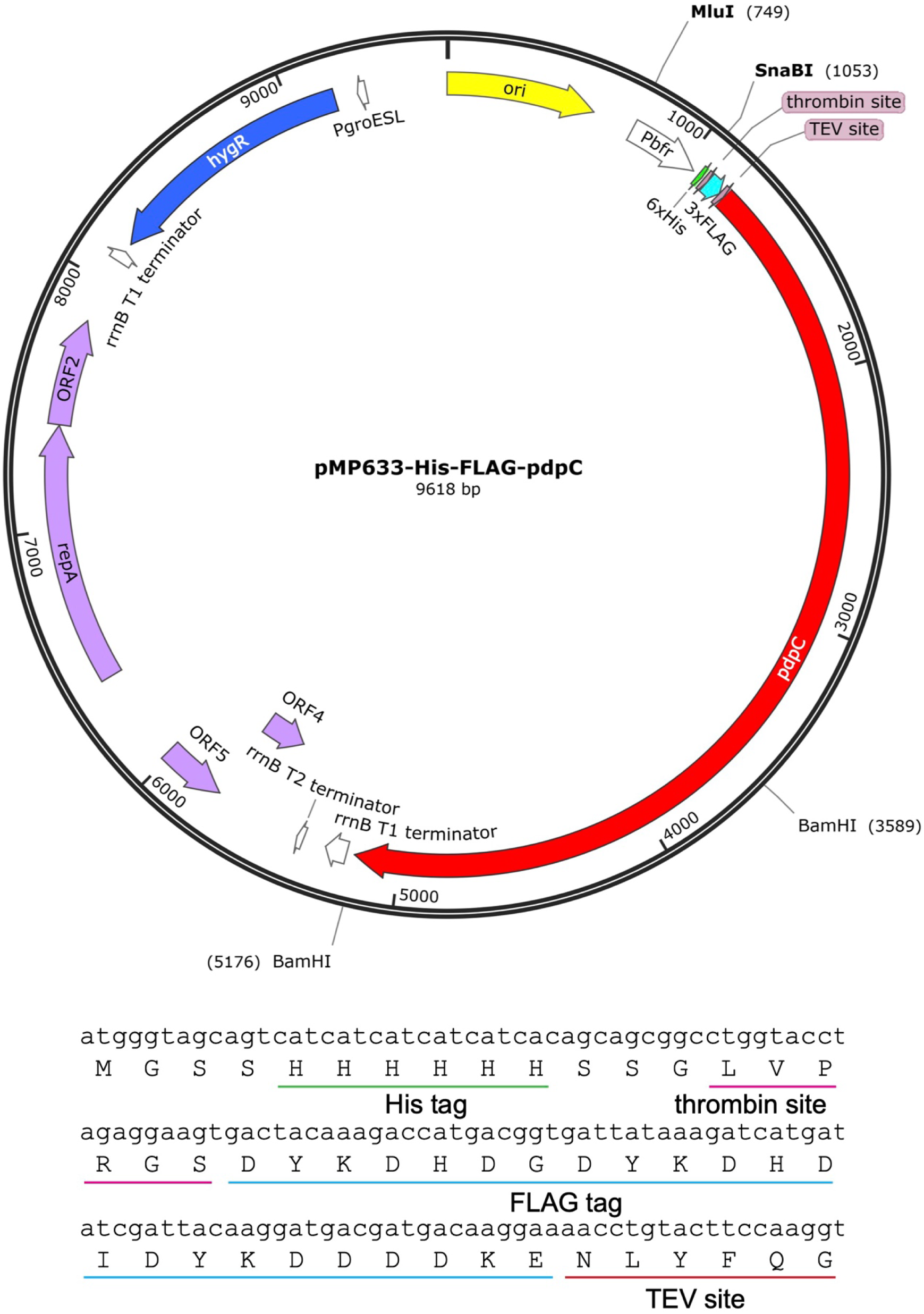
pMP633-Pbfr-His6-FLAG-pdpC plasmid map

**Supplementary Fig. S2:**
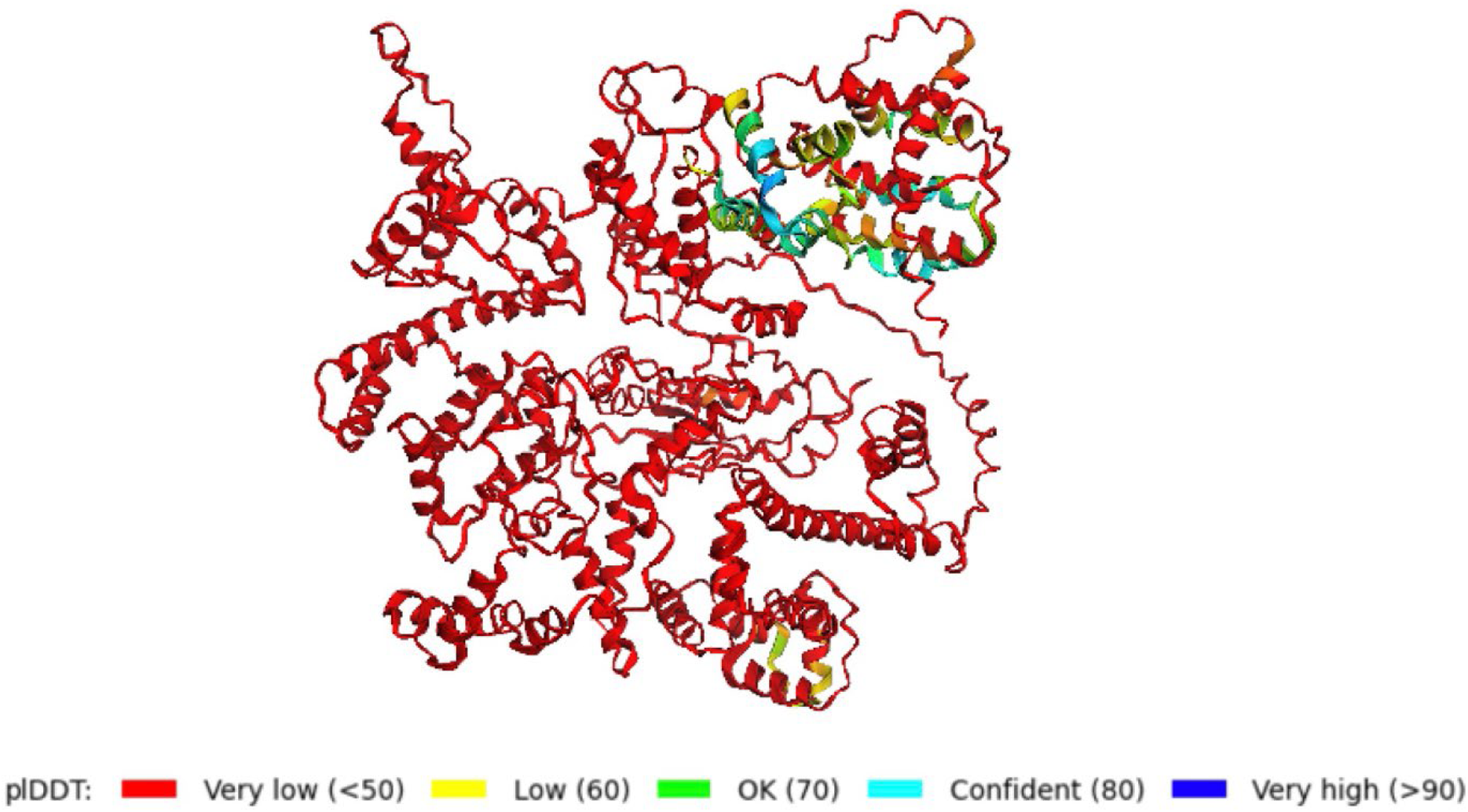
AlphaFold prediction of PdpC, colored by pLDDT. The prediction shows low confidence and poor correlation with the cryoEM structure determined in this study.

**Supplementary Fig. S3:**
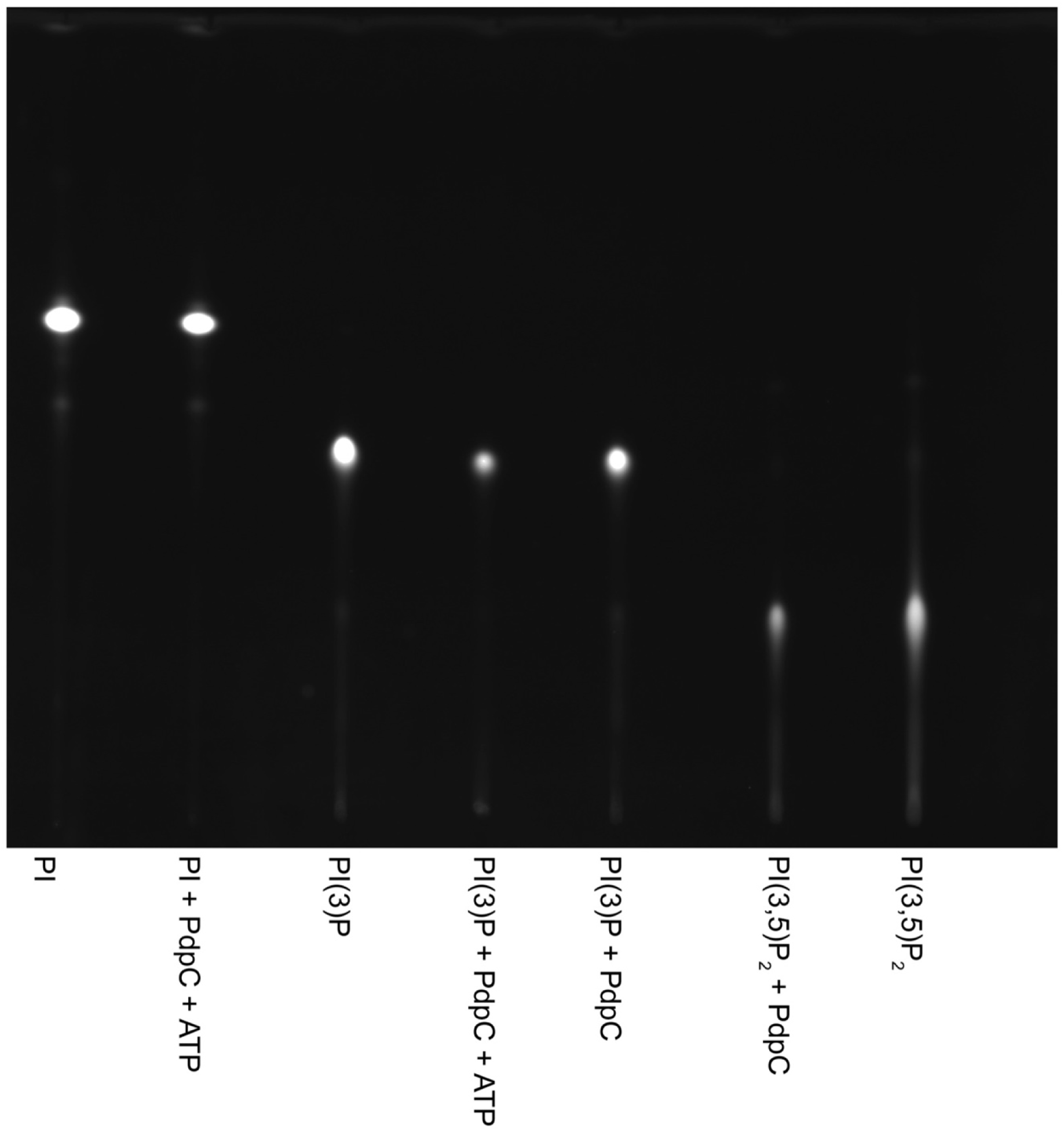
Phosphoinositide kinase and phosphatase activity are not detected in PdpC. PdpC was incubated with 50 µM BODIPY-labeled PI, PI(3)P, or PI(3,5)P2, with or without 10 mM ATP and 10 mM MgCl2 for 90 min at 37°C. The reaction was stopped by addition of acetone, dried under vacuum, and BODIPY-labeled lipids extracted with CHCl3, spotted onto a silica thin layer chromatography plate, developed in CHCl3:MeOH:Acetone:HOAc:Water (70:50:20:20:20), and imaged under blue light. Equal amounts of standard PI, PI(3)P, and PI(3,5)P2 that were incubated in the same buffer conditions without PdpC are included as reference controls.

**Table S1.** CryoEM data collection, refinement and validation statistics.

|  | PdpC<br>EMDB xxxx<br>PDB xxxx |
| --- | --- |
| **Data collection and processing** |  |
| Magnification | 81000 |
| Voltage (kV) | 300 |
| Electron exposure (e<sup>-</sup>/Å<sup>2</sup>) | 50 |
| Defocus range (μm) | -1.8 to -2.6 |
| Pixel size (Å) | 1.1 |
| Symmetry imposed | C1 |
| Particle number | 322627 |
| Map resolution | 3.4 |
| FSC threshold | 0.143 |
| **Refinement** |  |
| Map sharpening *B* factor (Å<sup>2</sup>) | -116 |
| **Model composition** |  |
| Non-hydrogen atoms | 7462 |
| Protein residues | 905 |
| Ligand |  |
| ***B* factors (Å<sup>2</sup>)** |  |
| Protein | 13.6 |
| Ligand |  |
| **R.m.s. deviations** |  |
| Bond lengths (Å) | 0.003 |
| Bond angle (°) | 0.690 |
| **Validation** |  |
| MolProbity score | 1.72 |
| Clashscore | 7.10 |
| Poor rotamers (%) | 0.24 |
| **Ramachandran plot** |  |
| Favored (%) | 95.29 |
| Allowed (%) | 4.71 |
| Disallowed (%) | 0 |

**Table S2.**
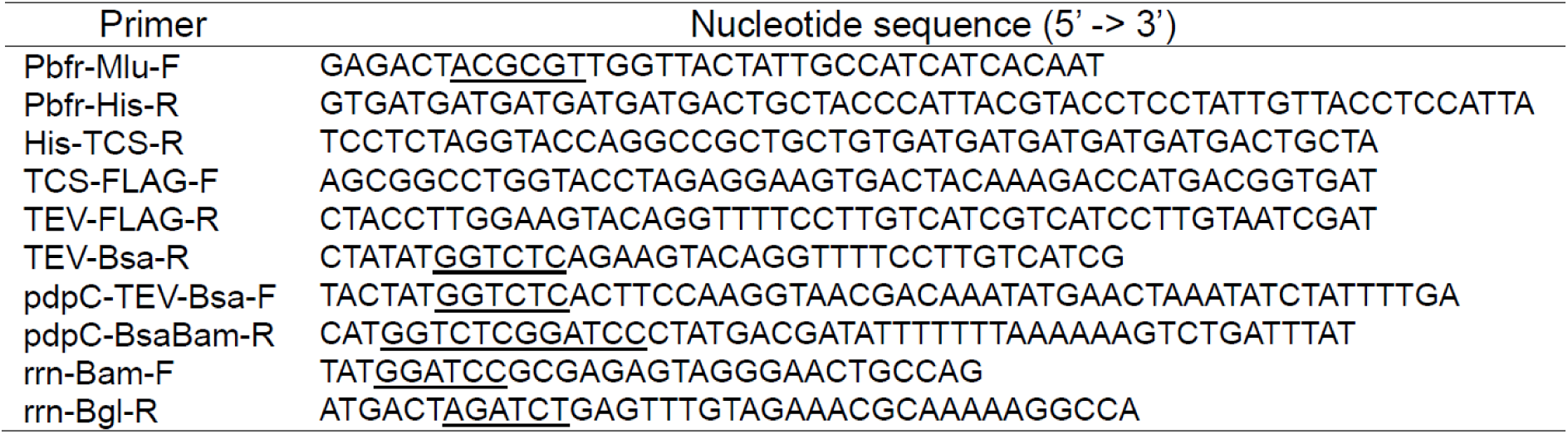
Oligonucleotide primers used in this study.

